# Virus-Like Particle Based-Vaccines Elicit Neutralizing Antibodies against the HIV-1 Fusion Peptide

**DOI:** 10.1101/2020.09.25.308957

**Authors:** Alemu Tekewe Mogus, Lihong Liu, Manxue Jia, Diane T. Ajayi, Kai Xu, Rui Kong, Jing Huang, Jian Yu, VRC Production Program, Peter D. Kwong, John R. Mascola, David D. Ho, Moriya Tsuji, Bryce Chackerian

**Author notes:** VRC Production Program: Nadia Amharref, Frank J. Arnold, Nathan Barefoot, Christopher Barry, Elizabeth Carey, Ria Caringal, Kevin Carlton, Naga Chalamalsetty, Anita Changela, Adam Charlton, Rajoshi Chaudhuri, Mingzhong Chen, Peifeng Chen, Nicole Cibelli, Jonathan W. Cooper, Hussain Dahodwala, Marianna Fleischman, Julia C. Frederick, Haley Fuller, Jason Gall, Isaac Godfroy, Daniel Gowetski, Krishana Gulla, Vera Ivleva, Lisa Kueltzo, Q. Paula Lei, Yile Li, Venkata Mangalampalli, Sarah O’Connell, Aakash Patel, Erwin Rosales-Zavala, Elizabeth Scheideman, Nicole A. Schneck, Zachary Schneiderman, Andrew Shaddeau, William Shadrick, Alison Vinitsky, Sara Witter, Yanhong Yang, and Yaqiu Zhang. Corresponding authors: (BC) and (MT).

## Abstract

Broadly neutralizing antibodies (bnAbs) isolated from HIV-infected individuals delineate vulnerable sites on the HIV envelope glycoprotein that are potential vaccine targets. A linear epitope at the N-terminal region of the HIV-1 fusion peptide (FP8) is the primary target of VRC34.01, a bnAb that neutralizes ~50% of primary HIV isolates. FP8 has attracted attention as a potential HIV vaccine target because it is a simple linear epitope. Here, we used platform technologies based on RNA bacteriophage virus-like particles (VLPs) to develop multivalent vaccines targeting the FP8 epitope. We produced recombinant MS2 VLPs displaying the FP8 peptide and we chemically conjugated synthetic FP8 peptides to Qβ VLPs. Both recombinant and conjugated FP8-VLPs induced high titers of FP8-specific antibodies in mice. A heterologous prime-boost-boost regimen employing the two FP8-VLP vaccines and native envelope trimer was the most effective approach for eliciting HIV-1 neutralizing antibodies. Given the potent immunogenicity of VLP-based vaccines, this vaccination strategy – inspired by bnAb-guided epitope mapping, VLP bioengineering, and optimal prime-boost immunization strategies – may be an effective strategy for eliciting bnAb responses against HIV.

## Introduction

Human immunodeficiency virus type 1 (HIV-1) continues to impose a significant burden of disease worldwide. Despite some progress in animal models ^1–3^, candidate vaccines evaluated to date have not yet been successful in inducing immunity in humans that protects from HIV infection. There is widespread agreement that the development of a successful HIV-1 vaccine will be dependent on the ability to induce potent protective antibodies capable of neutralizing diverse HIV-1 isolates. These antibodies, termed broadly neutralizing antibodies (bnAbs), are found in nearly 50% of HIV-infected individuals, but only after multiple rounds of immune selection and viral escape ^4,5^. These bnAbs commonly target a few sites of vulnerability on the HIV envelope glycoprotein (or Env trimer): the CD4 binding site ^5,6^, the trimer V1V2 apex ^7,8^, the variable glycan-V3 loop ^9,10^, the gp120-gp41 interface, the fusion peptide and the membrane-proximal external region ^11,12^. The isolation of bnAbs from HIV-infected patients and identification of their target epitopes on the conserved regions of HIV-1 Env trimer have provided possible pathways for vaccine design. However, many bnAbs recognize complex conformational epitopes or have undergone extensive affinity maturation from their germline state. These factors have complicated schemes for devising effective vaccine regimens for eliciting bnAb responses ^13^.

The HIV-1 fusion peptide is a critical component of the viral entry machinery that is composed of 15-20 hydrophobic residues at the N-terminus of the gp41 subunit of HIV-1 Env ^12^. A simple linear epitope consisting of 8 amino acids at the N-terminal region of the gp41 fusion peptide (called FP8) has been shown to be a primary target of a bnAb, VRC34.01, derived from an HIV-patient ^12,14^. Recently, Xu *et al*. designed an FP8-based vaccine in which the peptide is conjugated to keyhole limpet hemocyanin (KLH) through maleimide linkage chemistry ^3^. This FP8-KLH vaccine elicited fusion peptide-directed antibodies in mice, guinea pigs and rhesus macaques that were capable of neutralizing diverse strains of HIV-1 ^3^. Intriguingly, this FP8-KLH vaccine, in combination with multiple Env trimer boosts, elicited FP8-directed antibodies in rhesus macaques that neutralized 59% of 208 diverse viral strains ^15^.

Antibodies targeting FP8 that have been elicited by infection or by vaccination exhibit broad neutralization ^3,12,15^; however, neutralization activity against specific HIV-1 isolates is dependent on the structural diversity of the fusion peptide (FP), sensitivity to mutations at specific antibody interaction sites of the Env trimer outside of the FP and the natural diversity in FP8 sequences ^14,16^. FP, in the context of the native HIV Env trimer, adopts multiple conformations and orientations, which can both facilitate and complicate recognition of the FP8 epitope by bnAbs ^12,16,17^. X-ray crystallography and cryo-electron microscopy of the FP8-targeting antibodies in complex with the FP and Env trimer revealed diverse modes of antibody recognition of FP8 on the native HIV Env glycoprotein ^12,16^. Known FP8-targeting bnAbs approach the HIV Env trimer from diverse angles and recognize this epitope by penetrating through a highly conserved glycosylation site on gp120 (N88) ^16^. Mutations within this region of gp120 potentially affect the neutralization breadth of FP8-specific bnAbs ^14,16^. In addition, FP8 is not completely conserved in sequence across different subtypes of the HIV-1 Env ^12,14^. Functional mapping of infection- and vaccine-elicited antibodies against the FP has identified escape mutants and common variant sequences within the FP8 epitope ^14^. While functional mapping likely helps to re-center the protective domain on newly emerging HIV strains for subsequent rounds of immunogen design, the inherent structural diversity of FP may enable the design of conformationally diverse FP8 immunogens using multiple carrier protein or virus-like particle platforms, still capable of eliciting antibodies to recognize and neutralize FP in the context of native Env trimer.

Virus-like particles (VLPs) are a safe and highly immunogenic class of vaccines that have been utilized in many pre-clinical, clinical and post-clinical studies, with approved human vaccines available against hepatitis B virus ^18^, human papillomavirus ^19^ and hepatitis E virus ^20^. There has been increasing interest in bioengineering VLPs to serve as highly immunogenic platforms for surface display of foreign epitopes or antigens in a multivalent architecture ^21–24^. Bacteriophage VLPs are a particularly flexible and modular platform for vaccine development which allow the display of heterologous antigens on the surface of VLPs in a highly dense, repetitive array by several different techniques ^22,25^. For example, MS2 and PP7 bacteriophage coat protein single-chain dimers have been engineered for genetic insertion of heterologous peptides, and to produce *in vivo* assembled VLPs displaying heterologous peptides using bacterial cell factories ^26,27^. These recombinant VLPs are highly immunogenic and confer high immunogenicity to heterologous peptides displayed on their surfaces ^22,28^. In addition, bacteriophage VLPs can be chemically modified to display target antigens. For example, some bacteriophage VLPs, including Qβ, contain a high density of surface-exposed lysines which can be targeted for modification using various chemical techniques, enabling multivalent display of antigens on VLPs ^29^. Qβ VLPs are produced by recombinant expression of the Qβ coat protein in bacteria and by subsequent *in vivo* self-assembly of 180 monomers into VLPs. Target peptides which have been synthesized to contain a free terminal cysteine can then be conjugated to the VLP using an amine- and sulfhydryl-reactive bifunctional cross-linker. Qβ VLP based vaccines targeting antigens from both pathogens and self-antigens have been constructed and assessed in numerous preclinical studies ^27,30,31^ as well as human clinical trials ^32–34^.

In this study, we describe using both chemical conjugation and genetic insertion to construct microbially-synthesized RNA bacteriophage VLPs displaying the HIV-1 FP8 epitope. Both approaches enable multivalent display of the FP8 epitope on the surface of *Escherichia coli* (*E. coli*) produced bacteriophage VLPs. Both conjugated Qβ-FP8 and recombinant MS2-FP8 VLPs were recovered with high purity, and the recombinant VLPs maintained their *in vivo* assembly capacity. The FP8-VLPs were tested in different prime-boost regimens to elicit FP8-specific HIV-1 neutralizing antibody in mice. In particular, IgG isolated from mice immunized with an MS2-FP8 VLP prime, and boosts with Qβ-FP8 VLPs and native trimeric Env (BG505 DS-SOSIP), had the strongest neutralizing activity against prototype Clade A and B HIV-1 virus isolates. These studies suggest that a VLP-based vaccine could be a useful component of an FP8 targeted vaccine strategy for eliciting bnAbs against HIV-1.

## Results and Discussion

### Engineering and Characterization of FP8-displaying VLPs

Modification of the exterior facets of bacteriophage VLPs by genetic insertion or chemical conjugation techniques has enabled multivalent display of diverse heterologous epitopes on the surface of the VLPs ^21^. We previously showed that an engineered version of the MS2 bacteriophage coat protein, called the single-chain dimer, can be used to display target peptides on the surface of VLPs, either at a constrained loop (the AB-loop) ^26^ or at the N-terminus of the MS2 coat protein ^27,35^. FP8 is a short linear epitope, solvent accessible and recognized by bnAbs at the N-terminus of HIV-1 gp41 ^12,16^. Correspondingly, we chose to insert FP8 at the N-terminus of the MS2 single-chain dimer, reasoning that display of FP8 in this context would be most similar to its native conformation. In order to produce recombinant MS2-FP8 VLPs, the FP8 sequence was genetically inserted at the N-terminus of the single-chain dimer of MS2 coat protein (shown schematically in Fig. 1b). The recombinant VLP comprises 90 single-chain dimers of MS2 coat protein which self-assemble into FP8-displaying MS2 VLP *in vivo* in *E. coli*. Thus, recombinant MS2-FP8 VLPs are predicted to display exactly 90 copies of the FP8 peptide per particle.

**Fig. 1:**
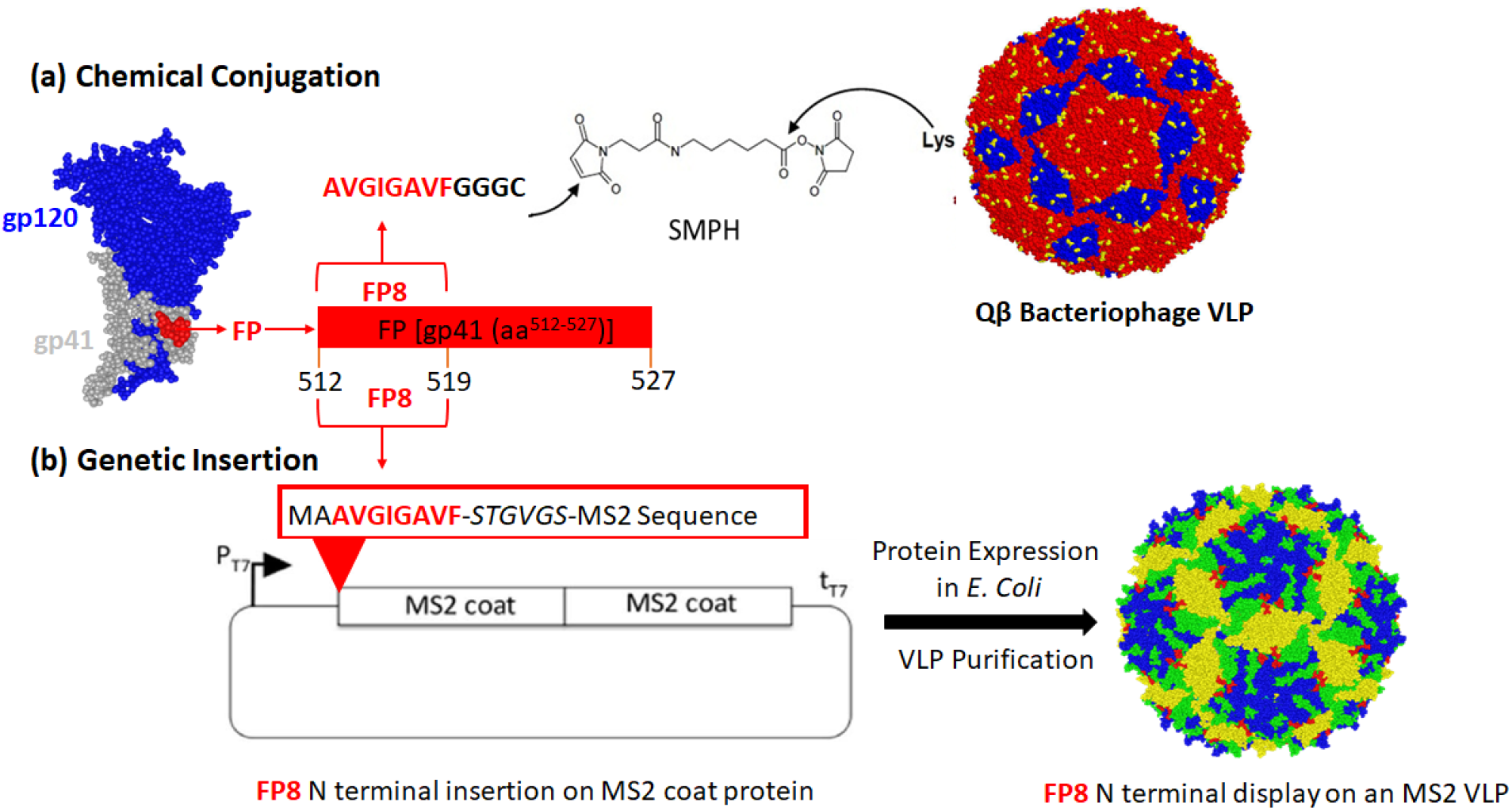
Strategy for displaying the HIV-1 fusion peptide epitope (FP8) on bacteriophage VLP (a) by chemical conjugation or (b) by constructing recombinant VLPs. In (a), the FP8 peptid was synthesized to contain a linker sequence with a terminal cysteine residue (−GGGC). The peptide was then conjugated to the surface-exposed lysine (Lys, in yellow) residues on microbially synthesized and *in vivo* assembled Qß bacteriophage VLPs (AB dimers in red, and CC dimers in blue) using a bifunctional crosslinker (SMPH). In (b), DNA sequence encoding the FP8 peptide was inserted into the N-terminus of MS2 bacteriophage coat protein single-chain dimer sequence on an expression vector. Recombinant MS2 VLPs displaying FP8 peptides were expressed from a plasmid in *E. coli*. The green, blue and yellow colors on MS2 VLP structure in the figure represent the residues derived from A-, B- and C-chains, respectively. The red color indicates the location of the N-terminal insertion site.

Microbially synthesized Qβ bacteriophage VLPs have been previously modified to display diverse antigens using a chemical crosslinker, in which peptides can be attached to exposed lysine residues that are abundantly displayed on the VLP surface ^22,36^. Fig. 1a illustrates this chemical technique for the construction of an FP8-displaying Qβ-VLP. The 8 amino acid FP8 epitope was chemically synthesized to include a C-terminal conjugation tag consisting of three glycines and a cysteine (FP8-*GGGC*). The synthetic FP8 peptides containing free cysteines were coupled to Qβ-VLPs using a bifunctional cross-linker (SMPH), yielding conjugated particles (Qß-FP8 VLPs). An analysis of conjugation efficiency by SDS-PAGE indicated that this technique resulted in an average of >200 FP8 peptides displayed per Qβ-VLP (Fig. 2a, Lane 3).

**Fig. 2:**
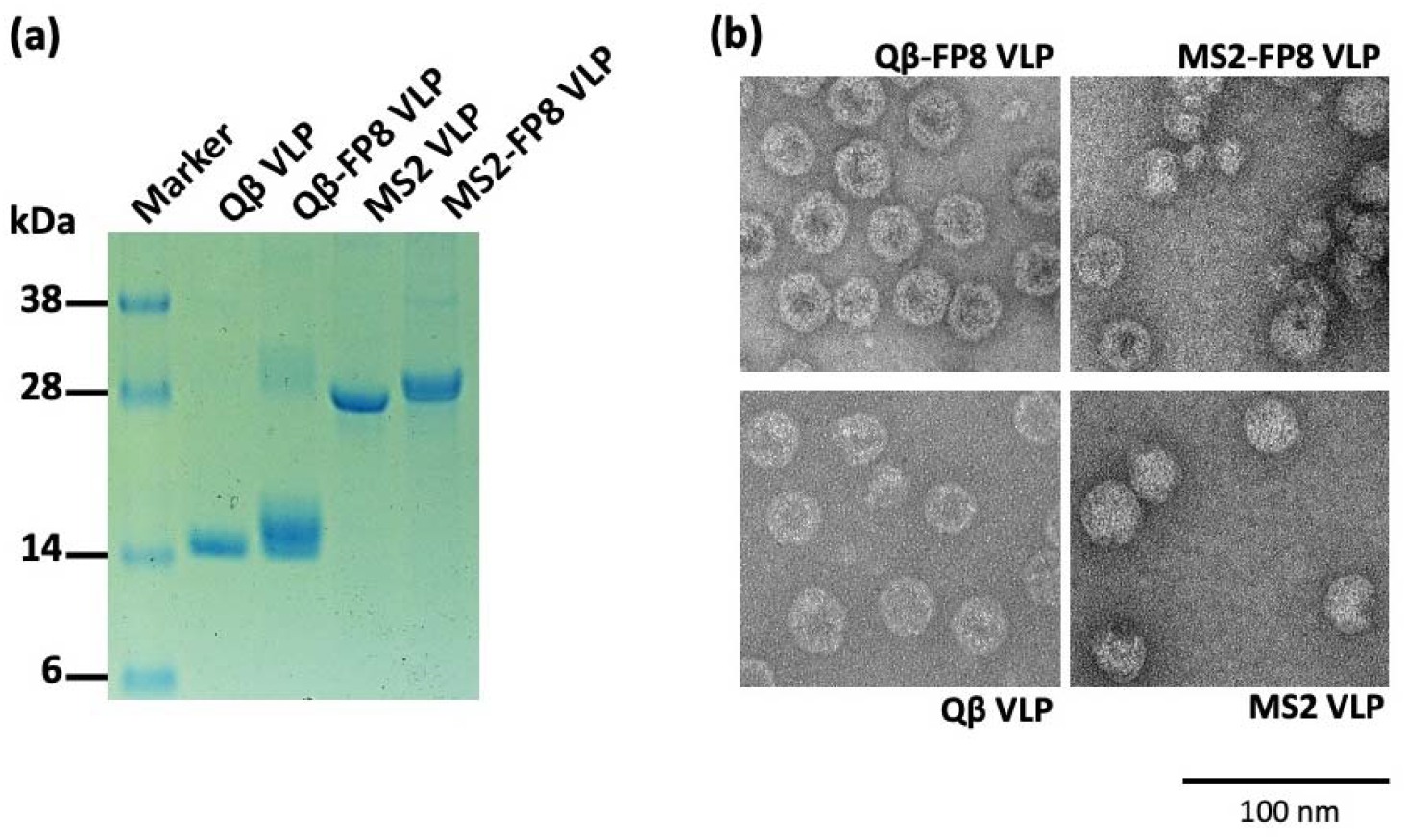
Characterization of conjugated (Qβ-FP8) and recombinant (MS2-FP8) VLPs. (a) SDS-PAGE analysis of proteins following the conjugation of FP8 peptides to microbially synthesized Qβ bacteriophage VLPs and the downstream processing of an *E. coli* expressed and *in vivo* assembled MS2-FP8 VLPs. The dominant bands on the gel correspond to product of the expected size. The ladder of bands in the Qß-FP8 VLP lane reflect individual copies of coat protein modified with 0, 1, 2, or more copies of the FP8 peptide. (b) Transmission electron micrograms (TEM) of conjugated (Qβ-FP8) and recombinant (MS2-FP8) VLPs in comparison to unconjugated and unmodified microbially synthesized Qβ and MS2 VLPs, respectively. Scale bar is 200 nm.

Fig. 2a shows SDS-PAGE analysis of purified Qβ-FP8 and MS2-FP8 VLPs. The lanes in Fig. 2a demonstrate successful conjugation of FP8 to the Qβ coat protein (Lane 3) and insertion of FP8 at the N-terminus of the single-chain dimer of MS2 coat protein (Lane 5). As shown by SDS-PAGE analysis (Fig. 2a), conjugated Qβ-FP8 VLPs and recombinant purified MS2-FP8 VLP were greater than 90% in purity. Centrifugation of conjugation reaction products using ultrafilters removed excess SMPH crosslinkers and FP8 peptides, yielding pure Qβ-FP8 VLPs. Selective salting-out precipitation of target proteins from *E. coli* cell lysates, with a polishing SEC step, resulted in the separation of recombinant *in vivo* assembled MS2-FP8 VLPs from most host cell protein contaminants.

To further characterize the FP8-displaying VLPs, we visualized VLPs by transmission electron microscopy (TEM) and also assessed VRC34.01 binding by ELISA. TEM micrographs of the conjugated (Qβ-FP8), the recombinant (MS2-FP8) and the unmodified (Qβ and MS2) VLPs are shown in Fig. 2b. Under TEM, Qβ-FP8 and MS2-FP8 VLPs were similar in morphology to the corresponding unmodified (Qβ and MS2) VLPs, confirming the particulate and multivalent nature of the FP8-displaying VLPs. Strong binding reactivity of Qβ-FP8 VLP and MS2-FP8 VLP with VRC34.01 (Fig. 3) using direct ELISA confirmed display of the FP8 epitope on the surface of VLPs. The stronger binding reactivity of Qβ-FP8 VLP is likely due to more numbers of FP8 peptides (>200 in Qβ VLP) in comparison to 90 FP8 peptides on the surface of each MS2 VLP. Thus, the peptide insertion and conjugation strategies were successful approaches to construct FP8-displaying MS2 and Qβ VLPs.

**Fig. 3:**
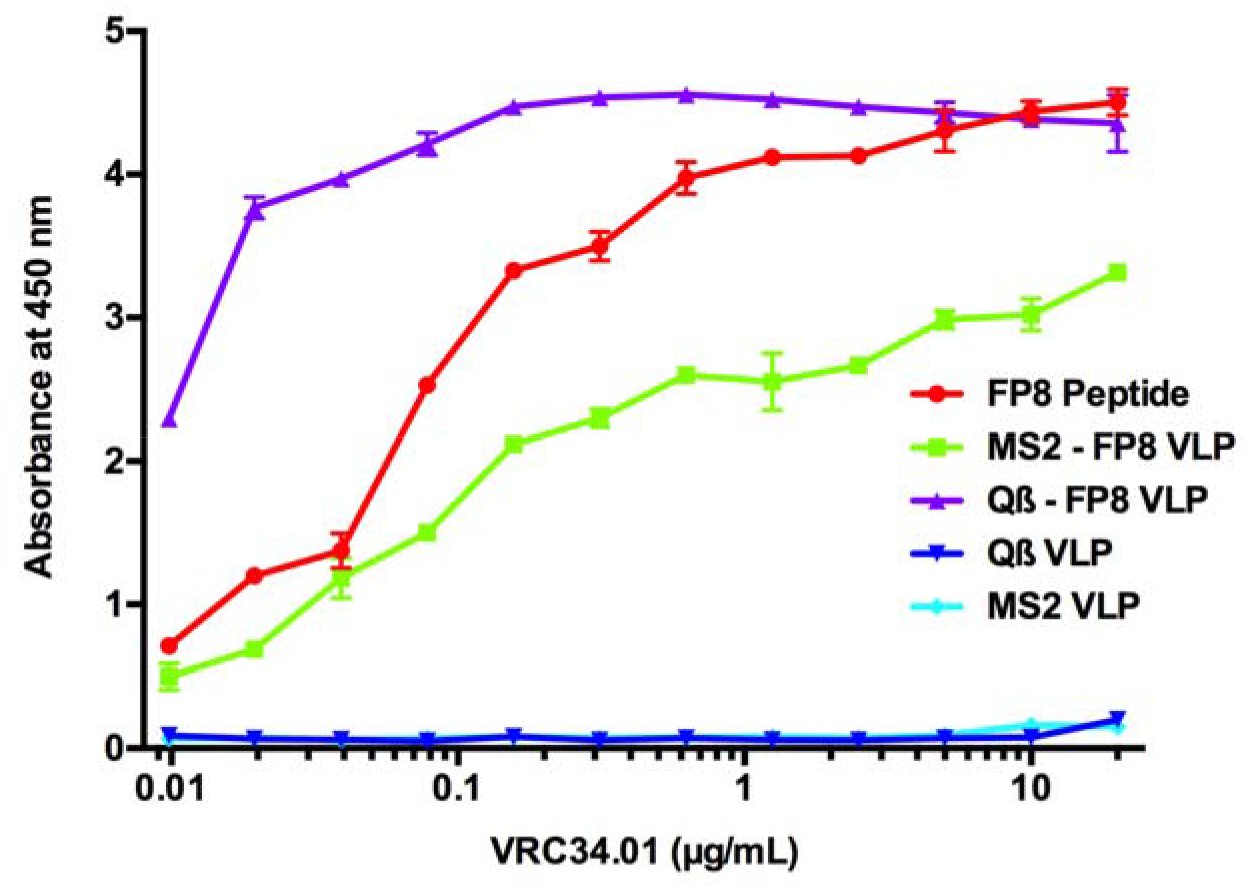
The FP8 peptide epitope is displayed on Qβ-VLPs and MS2-VLPs, as shown by ELISA. Qβ-FP8 VLP, MS2-FP8 VLP, wild-type (Qβ and MS2) VLPs (negative control), or FP8 peptide (positive control) were plated and probed with 2-fold dilutions of VRC34.01 bnAb. Reactivity of VRC34.01 confirms display of FP8 peptides on the surface of both VLPs.

### Immunogenicity of FP8-VLPs

Much effort has been directed to design immunogens and to find the optimal immunization strategies in order to efficiently elicit HIV neutralizing antibodies against discrete epitopes. The RV144 vaccine clinical trial, which showed modest protection from HIV infection, employed a prime/boost-based vaccination design with multiple immunizations ^37^. Other prime-boost vaccination approaches have shown potential in preclinical and clinical development of HIV vaccines ^38^. However, the ability to successfully elicit neutralizing antibodies is likely dependent on multiple factors, including the choice and order of priming-and boosting-immunogens. Here, we compared the immunogenicity of FP8-VLP vaccines in mice using both homologous and heterologous prime-boost regimens. Some groups of mice were also boosted with full-length trimeric Env protein (BG505 DS-SOSIP), because previous studies have shown that boosts with this antigen can amplify FP8-targeted antibody responses ^15^. As controls, mice were immunized with wild-type VLPs.

Fig. 4a outlines the immunogens used at prime, first boost and second boost in each of seven groups of mice. Mice sera were obtained after the first and second boosts and antibodies against the FP8 peptide and full-length BG505 DS-SOSIP were measured by ELISA. First, we measured the FP8- and BG505 DS-SOSIP-specific IgG titers following two immunizations with MS2-FP8 VLP (Groups I & II), Qβ-FP8 VLP (Groups IV & V) and the heterologous prime-boost groups (Groups III & VI). All of the vaccinated groups produced high titer IgG which recognized the FP8 peptide (Fig. 4b). The heterologous prime-boost regimen of FP8-VLPs (Group III & VI) induced a mean anti-FP8 antibody endpoint titer of more than 10^4^ after the first boost, significantly higher than the homologous prime-boost immunizations with MS2-FP8 VLPs (more than 13-fold higher; p < 0.001) or Qβ-FP8 VLPs (more than 3-fold higher; p < 0.05) [Fig. 4b]. In addition, two doses of Qβ-FP8 VLPs induced anti-FP8 antibody levels that were significantly higher than the groups immunized with MS2-FP8 VLPs (p < 0.05), indicating that increase antibody titers are likely due to the higher density of FP8 peptides (>200 versus 90) on the surface of Qβ VLPs. Anti-FP8 antibodies elicited by each of the experimental groups were also capable of recognizing full-length BG505 DS-SOSIP trimers (shown in Fig. 4c); similar to the FP8-specific responses, the groups that received a heterologous prime-boost regimen had the highest BG505 DS-SOSIP-specific IgG titers. Several other studies have shown that heterologous immunizations can be more effective than homologous immunizations ^38,39^. The specific mechanisms for the improved efficacy of the heterologous prime-boost vaccination are unknown, but it is possible that using multiple VLP platforms reduces the possibility that antibodies against the platform could interfere with responses against the FP8 peptide, mitigating the phenomenon referred to as carrier-mediated suppression ^40^. However, there is no evidence that carrier-mediated suppression reduces antibody responses to antigens displayed on VLPs; for example, suppression has not been observed in human clinical trials of Qβ bacteriophage VLP-based vaccines ^32–34,41^. Nevertheless, the present study demonstrates the effectiveness of a prime-boost approach using heterologous non-cross-reactive bacteriophage VLP carriers (Qβ and MS2) for enhancing FP8-specific antibody responses.

**Fig. 4:**
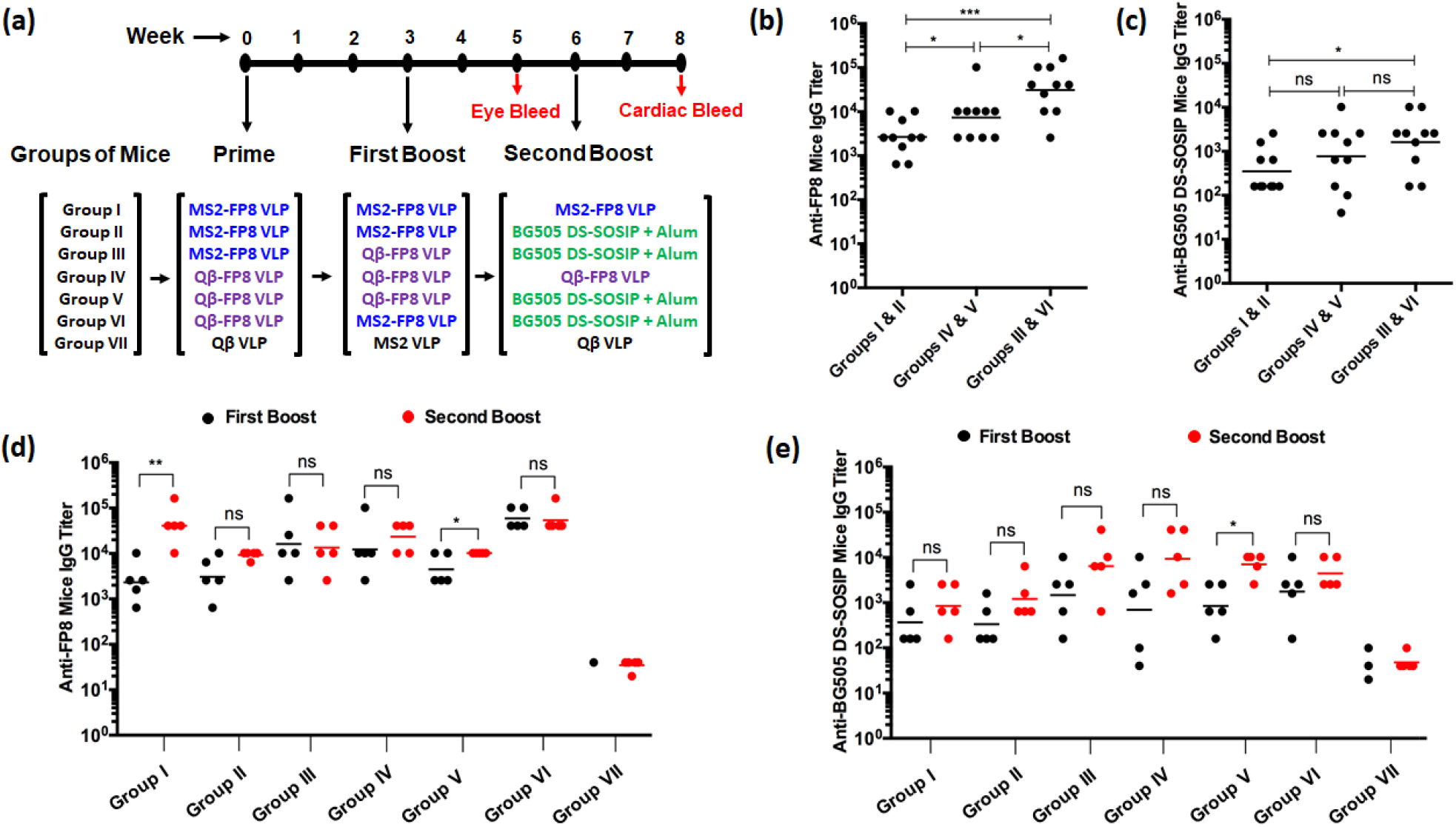
Immune responses in mice vaccinated with FP8-VLPs. (a) Immunization regimen. Groups of 5 mice were given three immunizations with FP8-VLPs (Groups I-VI) or control VLPs (Group VII). Mice were immunized at weeks 0 (prime), 3 (first boost), and 6 (second boost). At the second boost, some groups were immunized with native HIV-1 trimer (BG505 DS-SOSIP) plus Alum adjuvant. Serum collections occurred two weeks after the first boost [week 5] and the second boost [week 8]. Evaluation of the effect of homologous vs heterologous prime-boost regimen on the level of (b) FP8- and (c) BG505 DS-SOSIP-specific total IgG titers, which were measured by ELISA using week 5 immune sera. Data were combined for each MS2-FP8VLP, Qβ-FP8 VLP homologous-, and MS2-FP8 VLP/Qβ-FP8 VLP heterologous-prime boosting immunizations. Each dot represents the IgG titer from a single mouse. Lines represent geometric means (n = 10 per group). P values were calculated by unpaired two-tailed t test. *, P < 0.05, **, P < 0.01, ***, P < 0.001. (d) Anti-FP8 and (e) Anti-BG505 DS-SOSIP IgG end-point dilution antibody titers after the first boost (black dots) and second boost (red dots) were measured by ELISA analysis of sera. Lines represent geometric means (n = 5 per group). Statistical analysis of antibody titers after the first and second boost is presented. ^*^ P < 0.05 and ^**^ P < 0.01 indicate that there is a significant difference between the antibody titers of each group after the first and second boost, *ns* indicates no significant difference.

Although VRC34.01 recognizes a linear epitope, glycosylation events in envelope outside of the FP8 sequence can modulate antibody binding ^12,16^. Other FP8-based vaccine studies have demonstrated that boosting with native envelope can fine-tune the antibody responses and result in enhanced neutralization breadth of FP8-elicited antibodies ^3,14–17^. To assess the impact of boosting with trimer, some FP8-VLP immunized groups (shown in Fig. 4a) received an additional boost with adjuvanted BG505 DS-SOSIP. Antibody responses against FP8 (Fig. 4d, red dots) and BG505 DS-SOSIP (Fig. 4e, red dots) were measured by ELISA to compare antibody titers prior to and following the second boost. All of the groups immunized with FP8 vaccines produced high titer anti-FP8 (p < 0.0001) and anti-SOSIP (p < 0.01) IgG in comparison to the negative control (Group VII). Boosting with BG505 DS-SOSIP did not significantly increase the level of anti-FP8 antibody titer in comparison to its level after the first boost (Fig. 4d, red dots vs black dots, groups II, III, & VI). Boosting with BG505 DS-SOSIP did, however, result in an increase in anti-BG505 DS-SOSIP antibody titers. Anti-FP8 antibodies elicited by the groups of mice that received three homologous boosts of MS2-FP8 or Qβ-FP8 VLPs (groups I and IV) were also capable of binding to BG505 DS-SOSIP.

### FP8-VLPs elicit HIV-1 neutralizing antibodies

Production of high levels of FP8-specific antibodies in mice immunized with FP8-VLPs, and their binding reactivity to BG505 DS-SOSIP, are potentially indicative of FP8-specific neutralization activity against the virus. The ability of the anti-FP8 peptide sera to prevent viral infection *in vitro* was initially examined by a TZM-bl virus neutralization assay, using pooled sera from each group of mice against the Clade A primary HIV-1 isolate Q23.17. We chose Q23.17 because this virus is sensitive to neutralization by the positive control mAb, VRC34.01 ^3^. As shown in Fig. 5, sera from several of the groups of FP8-VLP immunized mice could neutralize HIV-1 Q23.17. The anti-FP8 sera from Group II and Group III mice were able to inhibit infection by 50% at a 1:80 dilution, and Group VI mice sera showed 50% neutralization at a 1:40 dilution (Fig. 5a). Immune sera obtained from Group I, Group IV, and Group V mice had weak neutralizing activity, but neutralization did not reach the 50% threshold at the lowest dilution tested.

**Fig. 5:**
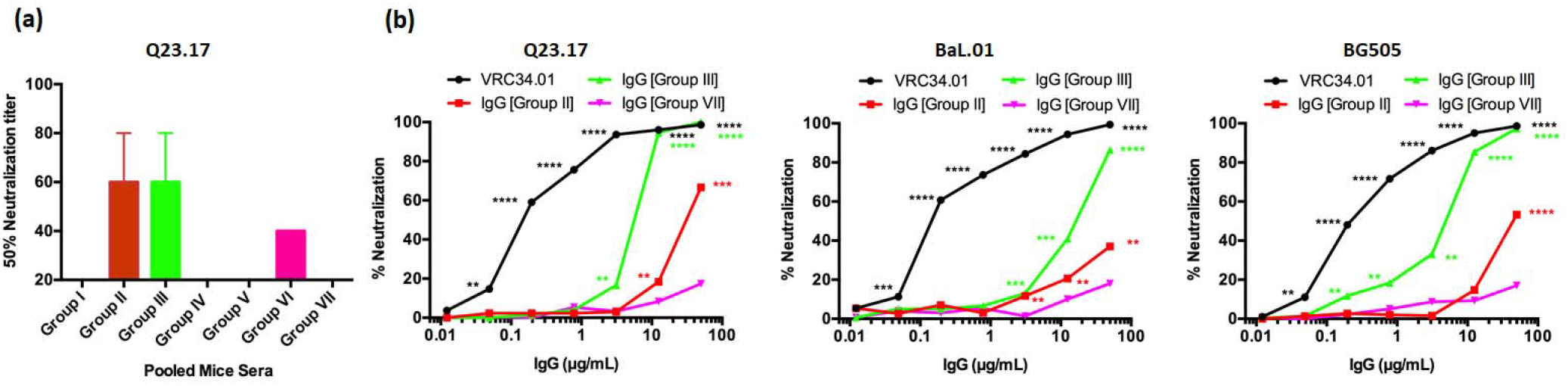
Neutralization activity of sera or purified IgG from FP8-VLP immunized mice. *In vitro* neutralization activity was determined using TZM-bl cells infected with Q23.17 (Clade A), BG505 (Clade A) and BaL.01 (Clade B) HIV-1 virus. (a) Neutralizing activity of pooled week 8 sera obtained from BALB/c mice immunized with various prime-boost-boost compositions containing FP8-VLPs alone or FP8-VLPs and BG505 DS-SOSIP was determined against Q23.17. The neutralization titer is expressed as the reciprocal of the highest serum dilution reducing cell infection by 50%. The lowest dilution tested was 1:20. (b) The viral neutralization activity of dilutions of purified serum IgG obtained from FP8-VLPs immunized mice with different prime boosting regimens was determined against Q23.17, BG505 and BaL.01, and is presented as percentage neutralization. VRC34.01 (positive) and IgG from Qβ/MS2 VLP immunized mice (negative) were used as controls. P values were calculated by paired two-tailed t test. *, P < 0.05, **, P < 0.01, ***, P < 0.001, ****, P < 0.0001.

Next, we tested the potency of purified IgG from the two groups [Group II & Group III] of mice with the strongest neutralizing activity against Q23.17 against a panel of three HIV-1 isolates. As shown in Fig. 5b, purified serum IgGs could neutralize prototype clade A (Q23.17 & BG505) and clade B (BaL.01) viruses. In particular, IgG isolated from Group III mice, which received a heterologous prime/boost/boost regimen, had potent neutralizing activity, exhibiting nearly complete neutralization of isolates Q23.17 and BG505 at a concentration of 12.5 μg/mL (Fig. 5b). Purified IgG from Group II also neutralized HIV-1, albeit with lower potency. Taken together, these data highlight the importance of heterologous boosting in generating FP8-specific antibodies with HIV-1 neutralizing activity.

Interestingly, sera from some immunized groups with high anti-FP8 antibody levels (such as in Group I) or high titer antibodies against BG505 DS-SOSIP of (such as in Group IV) did not neutralize HIV-1. The lack of correlation between the high level of anti-FP8 titers and the elicited neutralization activity of the mice sera suggests a critical parameter may be recognition of a specific conformation of the peptide, or, perhaps, appropriate affinity maturation of the responses, but most likely not the overall titer of reactive antibodies. Affinity maturation often increases the affinity, avidity and neutralization activity of antibodies through multiple rounds of somatic hypermutation and selection in the germinal center ^42–44^. Recent studies have used innovative vaccine design approaches and immunization strategies to trigger B cell precursors expressing germline receptors and then direct affinity maturation toward HIV-1 broad neutralizing antibodies ^1,44–46^. Xu *et al*., ^3^ demonstrated that an immunogen design based on FP8 linked to KLH and an immunization strategy, involving FP8-KLH priming and Env-trimer boosting, elicited FP8-directed cross-clade neutralizing antibodies in mice, guinea pigs and non-human primates. Multiple FP8-KLH primes and Env-trimer boosts increased FP8-directed cross-clade HIV neutralization breadth ^14,47^. These published results ^3,15,47^ and the results described in this study highlight the importance of the prime/boost compositions and immunization strategies and indicate that a sequence of immunizations with different immunogens may be a key to guide affinity maturation of FP8-directed bnAbs. In addition, well validated genetic and structural approaches for identification and characterization of neutralizing antibody lineages in vaccinated animals ^3,14^, open the opportunity to find an optimal immunogen design and immunization strategy that can induce the broadest FP8-directed neutralization.

### Conclusions

FP8, a site of vulnerability on the HIV-1 Env glycoprotein and a target for infection-elicited bnAbs, has become a promising target for HIV-1 vaccine design. Our aim was to develop new HIV vaccine candidates by displaying the FP8 peptide on the surface of microbially synthesized RNA bacteriophage VLPs and to find an optimal prime-boost immunization strategy for eliciting high titer FP8-directed HIV-1 neutralizing antibodies. We used two techniques, chemical conjugation and genetic insertion, to produce bacteriophage VLPs which display the FP8 peptide in a multivalent fashion. Both approaches yielded FP8-displaying VLPs which reacted strongly with an FP8-binding bnAb, VRC34.01, and, in mice, elicited antibodies that bound to native Env trimer. The FP8-VLPs were tested in different prime-boost regimens to elicit FP8-specific HIV-1 neutralization activity in mice. Immunization with MS2-FP8 VLP prime, Qβ-FP8 VLP first boost and alum adjuvanted BG505 DS-SOSIP second boost induced the most potent HIV-1 neutralizing antibody responses, highlighting the importance of a heterologous boosting regimen. The results in this study demonstrate that an approach combining the peptide display platform based on the RNA bacteriophage VLPs and a prime-boost-boost immunization strategy allowed the elicitation of FP8-specific HIV-1 neutralizing antibody in mice. These promising results warrant further studies to explore the full potential of FP8-VLP vaccine designs with multiple prime-boost compositions for the elicitation of the broadest FP8-directed neutralizing antibody responses. Further study is needed to identify and characterize FP8-directed broad neutralizing antibody lineages in vaccinated animals as well as to explore different adjuvants and multiple Env trimer boosts to increase neutralization titer and breadth.

## Materials and methods

### Construction of FP8-displaying recombinant VLPs

Plasmid pDSP62, which encodes the single-chain dimer version of the MS2 bacteriophage coat protein, was generated previously ^48^. The gene fragment encoding FP8 with as well as sequence encoding a flanking STGVGS linker sequence was cloned at the 5’ end of the single-chain dimer sequence by PCR. Briefly, a forward PCR primer (5' GCGCCATGGCAGCGGTTGGCATTGGAGCAGTTTTCTCAACCGGAGTTGGAAGCGCAAGCAATTTCACGCAATTTG 3') was designed to contain nucleotide sequence encoding a *NcoI* restriction site, a start codon and an alanine amino acid, FP8 sequence, a linker sequence and a sequence that is complementary to the N-terminus of the MS2 single-chain dimer coat protein. The reverse primer E3.2 (5' CGGGCTTTGTTAGCAGCCGG 3'), that anneals downstream of a unique *BamHI* site in the pDSP62 plasmid, was described previously ^27^. The primers were used to amplify a gene fragment using plasmid pDSP62 as a PCR template DNA. Amplified MS2-FP8 PCR fragment was digested with *NcoI* and *BamHI* restriction enzymes and cloned into pDSP62 plasmid using these restriction sites. The cloned construct was sequenced to confirm the presence of the FP8 peptide insert and designated as pDSP62-FP8.

### Production and purification of FP8-displaying recombinant VLPs

pDSP62-FP8 plasmids were transformed into C41 *E. coli* cells by electroporation. Transformed C41 cells were grown at 37° C using Luria Bertani broth containing 60 μg/mL kanamycin until the cells reached an OD600 of 0.6. MS2-FP8 protein expression was induced using 0.4 mM isopropyl-β-D-1-thiogalactopyranoside and grown at 37° C overnight. Cell pellets were collected and re-suspended using a lysis buffer [50 mM Tris-HCL, 100 mM NaCl, 10 mM ethylenediaminetetraacetic acid, pH 8.5]. Cells were lysed by sonication and cell lysates were clarified by centrifugation (15000g, 20 min, 4° C). Soluble MS2-FP8 VLPs were purified by selective-salting out precipitation using 70% saturated (NH_4_)_2_SO_4_, followed by an additional polishing size exclusion chromatography (SEC) step using a Sepharose CL-4B column. The column was pre-equilibrated with a purification buffer (40 mM Tris-HCl, 400 mM NaCl, 8.2 mM MgSO_4_, pH 7.4). MS2-FP8 VLPs were concentrated from SEC purified fractions by ultra-centrifugal filtration using Amicon^®^ Ultra-15.0mL 100 K membrane (Merck Millipore Ltd., Tullagreen, Carrigtwohill, Co. Cork, Ireland; 3000g, 10 min at 22° C).

### Conjugation of FP8 to Q β bacteriophage VLPs

Qβ-VLPs were produced in *E. coli* using methods as previously described to produce MS2 bacteriophage VLPs ^27^. The FP8 peptide (AVGIGAVF) was synthesized (GenScript) and modified to include a C-terminal cysteine residue preceded by a 3-glycine-spacer sequence. FP8-GGGC peptides were conjugated to Qβ-VLPs using the bifunctional cross-linker succinimidyl 6-[(β-maleimidopropionamido) hexanoate] (SMPH; Thermo Fisher Scientific) as described previously for the conjugation of amyloid-beta (Aβ) peptides ^49^.

### Characterization of FP8-displaying VLPs

Conjugated (Qβ-FP8) and recombinant (MS2-FP8) VLPs were run on a 10% sodium dodecyl sulfate polyacrylamide gel electrophoresis (SDS-PAGE) gel stained with Coomassie blue to check the efficiency of conjugation and the purity of the VLPs. All VLP concentrations were estimated from SDS-PAGE gel protein bands corresponding to Qβ, Qβ-FP8, MS2 and MS2-FP8 coat proteins using known amounts of hen egg lysozyme as the standard protein. Visualization of the VLPs with transmission electron microscope (TEM) was performed as previously reported ^50^. The display of FP8 peptides on the surface of Qβ and MS2 VLPs was confirmed by enzyme-linked immunosorbent assay (ELISA) as follows: the 96-well Immulon^®^ 2 plate (Thermo Fischer Scientific) was coated with 500 ng of each of Qβ-FP8 VLP, MS2-FP8 VLP, positive control (FP8 peptide) and negative controls (Qβ and MS2 VLPs) in duplicate diluted in 50 μL of phosphate-buffered saline (PBS) in each well. The plate was incubated at 4 ° C for overnight and blocked for 1 h at room temperature with blocking buffer (0.5% dry milk in PBS). Wells were washed two times with PBS and incubated for 2 h at room temperature with VRC34.01 initially at 50-fold dilutions followed by two-fold dilutions in blocking buffer. VRC34.01 was expressed by transient transfection from with plasmids coding for the heavy and light chains of the antibody as previously described ^12^. The wells were washed five times with PBS and probed with horseradish peroxidase (HRP)-conjugated secondary antibody [goat anti-human IgG (Jackson ImmunoResearch; 1:5000)] for 1 h. The reaction was developed using TMB (Thermo Fischer Scientific) and stopped using 1% HCl. Reactivity of VRC34.01 for the target FP8 peptides was determined by measuring optical density at 450 nm (OD450).

### Mice immunizations

All animal studies were performed in accordance with guidelines of the University of New Mexico Animal Care and Use Committee (Approved protocol #: 19-200870-HSC). Seven groups of five female Balb/c mice, aged 6-8 weeks, were obtained from Jackson Laboratory. All groups of mice received prime, first and second booster doses intramuscularly at three-week intervals. VLPs were immunized at a dose of 5 μg without exogenous adjuvant. At the second boost, some groups were immunized with 25 μg of BG505 DS-SOSIP.664 and formulated with Alhydrogel^®^ (InvivoGen, USA) at a 1:1 (volume: volume) ratio. BG505 DS-SOSIP.664 was produced from a CHO-DG44 stable cell line, purified using non-affinity chromatography and was antigenically similar to trimers described previously ^51^. The injection volume was 50 μL for all formulations. Blood was collected two weeks following the first and second boosts; serum was isolated and stored at −20 ° C.

### Characterization of antibody responses

FP8- and BG505 DS-SOSIP-specific mice serum IgG titers were determined by end-point dilution ELISA. For FP8 peptide ELISAs, Immulon^®^ 2 plates (Thermo Scientific) were incubated with 500 ng streptavidin (Invitrogen) in PBS, pH 7.4, for 2 h at 37 ° C. Following washing, SMPH was added to wells at 1.0 μg/well and incubated for 1 h at room temperature. The peptide was added to the wells at 1.0 μg/well and incubated overnight at 4° C. BG505 DS-SOSIP ELISAs were performed based on a previously reported lectin captured trimer method, with slight modifications ^52^. The plates were coated with 50 μL/well of 2.0 μg/mL of Lectin (*Galanthus nivalis)* [Sigma Aldrich] in PBS overnight at 4°C. After blocking, 50 μL/well of 2.0 μg/mL BG505 DS-SOSIP in blocking buffer was added and incubated for 2 h at room temperature. For all ELISAs, plates were blocked with 0.5% milk in PBS (150 μL/well) for 1 h at room temperature, and four-fold dilutions of mice sera (starting at 1:40) were added to each well and incubated for 2 h. The wells were probed with horseradish peroxidase (HRP)-conjugated secondary antibody [goat anti-mouse-IgG (Jackson Immuno-Research; 1:5000)] for 1 h. The reaction was developed using TMB (Thermo Fischer Scientific) and stopped using 1% HCl. Reactivity of sera for the target antigen was determined by measuring optical density at 450 nm (OD_450_). Wells with twice the OD_450_ value of background were positive and the highest dilution with a positive value was considered as the end-point dilution titer.

### Immunoglobulin purification

Dynabeads™ Protein G (Invitrogen) were used to purify IgG from pooled mice sera from selected immunized groups (Groups II, III, and VII). First, the Dynabeads™ Protein G were washed with PBS by placing the tube in a DynaMag™ magnet (Thermo Fischer Scientific). The mice sera were incubated with the Dynabeads™ Protein G in the tubes for 40 minutes at room temperature. The Dynabeads were washed with PBS by placing the tubes in the DynaMag™ magnet to remove unbound materials. IgG was eluted using 0.2 M glycine/HCl buffer (pH 2.7) and then brought to neutral pH using 1 M Tris buffer (pH 9). The concentration of purified mouse IgG was measured by reading absorbance at 280 nm.

### HIV-1 neutralization assay

The neutralization activity of pooled sera obtained from Balb/c mice immunized with various prime-boost-boost compositions was determined by using the TZM-bl assay. Sera were assessed in duplicate at eight 4-fold dilutions, starting at a 1:20 dilution. Sera were incubated with HIV-1 Q23.17 (Clade A) virus prior to adding to TZM-bl cells and incubated for 3 days. After removing supernatant, the cells were lysed, and β-Gal was added to detect the β-galactosidase activity, and a 50% inhibitory concentration (IC_50_) were determined. The IC_50_ were expressed as the reciprocal of the serum dilutions required to inhibit infection by 50%. Serum IgGs were purified [section 2.7 above] from the immune sera (Group II and Group III mice) capable of inhibiting infection by 50%. Purified serum IgGs were tested against Q23.17 (Clade A), BG505 (Clade A) and BaL.01 (Clade B) HIV-1 virus isolates in triplicate at using seven 4-fold dilutions, starting at 50μg/mL, and data are reported as percent (%) neutralization inhibition. Purified IgG from Group VII mice and VRC34.01 bnAb were used as negative and positive controls, respectively.

### Statistical analysis

Statistical analysis was carried out using GraphPad Prism Version 6.00 (GraphPad Software Inc., CA, USA). Comparison between two groups was performed with paired t test or unpaired two-tailed t test. Groups were considered significantly different (**\***) at P < 0.05.

## Acknowledgements

The authors acknowledge research funding from National Institute of Health (NIH) [BC; R01AI124378]. Support for this work was also provided in part by the Intramural Research Program of the Vaccine Research Center, National Institute of Allergy and Infectious Diseases, National Institutes of Health (PDK and JRM).

